# The *Brassica napus* Wall-Associated Kinase-Like (WAKL) gene *Rlm9* provides race-specific blackleg resistance

**DOI:** 10.1101/815845

**Authors:** Nicholas J. Larkan, Lisong Ma, Parham Haddadi, Miles Buchwaldt, Isobel A. P. Parkin, Mohammad Djavaheri, M. Hossein Borhan

**Affiliations:** Armatus Genetics Inc., Saskatoon, SK, Canada; Agriculture & Agri-Food Canada, Saskatoon, SK, Canada

**Keywords:** *Leptosphaeria maculans*, blackleg, *Brassica napus*, *Rlm9*, *AvrLm5-9*, disease resistance, wall-associated kinase-like

## Abstract

In plants, race-specific defense against microbial pathogens is facilitated by resistance (*R*) genes which correspond to specific pathogen avirulence (*Avr*) genes. This study reports the cloning of a blackleg *R* gene from *Brassica napus* (canola); *Rlm9*, which encodes a wall-associated kinase-like (WAKL) protein, a newly-discovered class of race-specific plant RLK resistance genes. *Rlm9* provides race-specific resistance against isolates of *Leptosphaeria maculans* carrying the corresponding avirulence gene *AvrLm5-9*, representing only the second WAKL-type *R* gene described to date. The Rlm9 protein is predicted to be cell membrane-bound yet appears to have no direct interaction with AvrLm5-9. *Rlm9* forms part of a distinct evolutionary family of RLK proteins in *B. napus,* and while little is yet known about WAKL function, the *Brassica-Leptosphaeria* pathosystem may prove to be a model system by which the mechanism of fungal avirulence protein recognition by WAKL-type *R* genes can be determined.

## Introduction

Plants detect invading microbial pathogens through the perception of conserved pathogen-associated molecular patterns (PAMPs). Perception of PAMPs by the extracellular pattern recognition receptors (PRR), consisting of various receptor-like kinases (RLKs) and receptor-like proteins (RLPs), initiates host PAMP triggered immunity (PTI), the first layer of defense against pathogens. PRRs also respond to damage-associated molecular patterns (DAMPs); host peptides and oligosaccharide fragments such as pectin-derived oligogalacturonides (OGs) released during pathogen breaching of the plant cell wall (Yang *et al*., 2012; Zipfel *et al*., 2014; Boutrot & Zipfel, 2017). Pathogens have evolved to overcome PTI by secreting small proteins called effectors, which often target components of the plant’s defense pathways. In turn plants are armed with an array of highly diverse resistance (*R*) genes which encode both cytoplasmic and extracellular receptors to perceive pathogen effectors and trigger a rapid and robust immune response called effector-triggered immunity (ETI) (Bent & Mackey, 2008; Jones & Dangl, 2006; Stotz *et al*., 2014). Generally, PTI provides broad-spectrum resistance as it is triggered by conserved signals, such as bacterial flagellin, fungal chitin or host-derived OGs, while ETI is specific to the races of single species of pathogen which produces the matching effector(s). However, PTI and ETI share common signaling pathways (Katagiri & Tsuda, 2010) and the distinction between the two is blurred. There are examples of PRR proteins that confer race specificity and *R* genes that are now being considered as PRRs (Thomma et al., 2011; Rodriguez-Moreno et al., 2018).

Plant RLK proteins are a large and diverse gene family which underwent massive expansion after the divergence of the plant and animal lineages (Shiu & Bleecker, 2001). RLKs are defined by a common set of domains; a signal peptide, a single transmembrane domain and a cytoplasmic kinase domain. The extracellular regions of the proteins vary, adapted to the recognition of diverse signals. Based on their conserved intracellular kinase domains plant RLKs form a monophyletic group distinct from other eukaryotic kinases (Shiu & Bleecker, 2001). Membrane bound RLPs, which feature extracellular leucine-rich repeat (eLRR) domains involved in protein recognition but lack the intracellular kinase domain of RLKs, constitute a class of R proteins that confer resistance upon perception of apoplastic pathogen effectors (Stotz *et al*., 2014; Jamieson *et al*., 2018). We previously reported the cloning of two RLP type resistance genes, *LepR3* and *Rlm2*, residing in the A genome of the allotetraploid (AACC) *Brassica napus* (canola, rapeseed) (Larkan et al., 2013; Larkan et al., 2015), conferring resistance against races of the fungal pathogen *Leptosphaeria maculans* (Lm) with the matching effectors AvrLm1 and AvrLm2, respectively (Ghanbarnia *et al*., 2015; Gout *et al*., 2006). Both LepR3 and Rlm2 pair with the RLK SOBIR1 (Ma & Borhan, 2015; Larkan *et al*., 2015) as RLPs, lacking any intracellular kinase, require a partner to transmit a signal across the plasma membrane and to activate cytoplasmic signal-transduction cascades (Liebrand *et al*., 2014).

The *Brassica R* genes *Rlm3, Rlm4, Rlm7* and *Rlm9* confer race-specific resistance against blackleg disease caused by Lm. They form a tight genetic cluster on chromosome A07 and may possibly be allelic variants of the same *R* locus (Larkan *et al*., 2016). The corresponding avirulence (*Avr*) genes *AvrLm3, AvrLm4-7* and *AvrLm5-9* have been cloned from Lm and all encode small cysteine-rich secreted proteins (Parlange *et al*., 2009; Plissonneau *et al*., 2016; Ghanbarnia *et al*., 2018). Recognition of both AvrLm3 and AvrLm5-9 by Rlm3 and Rlm9, respectively, is masked in the presence of AvrLm4-7. However AvrLm4-7 neither interferes with the expression of, nor directly interacts with, AvrLm3 or AvrLm5-9, nor do AvrLm3 and AvrLm5-9 interact (Ghanbarnia *et al*., 2018). To investigate this complex system of pathogen recognition we have pursued the cloning of *Brassica R* genes from the *Rlm3/4/7/9* gene cluster. Here we report cloning of *Rlm9* from the *B. napus* cultivar ‘Darmor’ and show that it encodes a wall-associated kinase-like (WAKL) protein, a newly-emerging class of race-specific plant RLK resistance genes (Keller & Krattinger, 2018).

## Results

### *Rlm9* encodes a Wall-associated Kinase-like Protein

Through molecular mapping, the physical interval of the *Rlm9* locus had previously been defined as approximately 4.3 Mb of chromosome A07 (Larkan *et al*., 2016) of the *Brassica napus* reference genome ‘Darmor-*bzh’* (Chalhoub *et al*., 2014), an *Rlm9* variety. *Rlm9* is genetically clustered with the other blackleg *R* genes *Rlm3, Rlm4* and *Rlm7*, all of which were shown to co-segregate with the microsatellite marker sR7018 positioned at approximately 16 Mb on chromosome A07 (Larkan *et al*., 2016). Using this information along with previously generated genomic information for the *Rlm3* locus (Mayerhofer *et al*., 2002) we searched the physical interval of the *Rlm3-4-7-9* cluster on the *B. napus* Darmor-*bzh* (*Rlm9*) genome for genes with similarity to *R* genes and expression in response to pathogen infection. BnaA07g20220D, a wall associated kinase-like protein (WAKL) encoding gene was identified as the best candidate for *Rlm9*. The WAKL is predicted to encode a 794 aa protein with the typical features of the wall-associated kinase-like family of *Arabidopsis thaliana* (Verica & He, 2002), showing the highest homology to *A. thaliana* WAKL10 (At1g79680, 69% amino acid identity). The gene consists of three exons encoding a transmembrane receptor protein, which contains predicted extracellular domains for pectin and calcium binding (wall-associated receptor kinase galacturonan-binding (GUB_WAK) and epithelial growth factor (EGF)-like Ca^2+^ domains, respectively), a C-terminal WAK domain and an intracellular serine/threonine protein kinase domain with a guanylyl cyclase motif (Figure 1).

**Figure 1.**
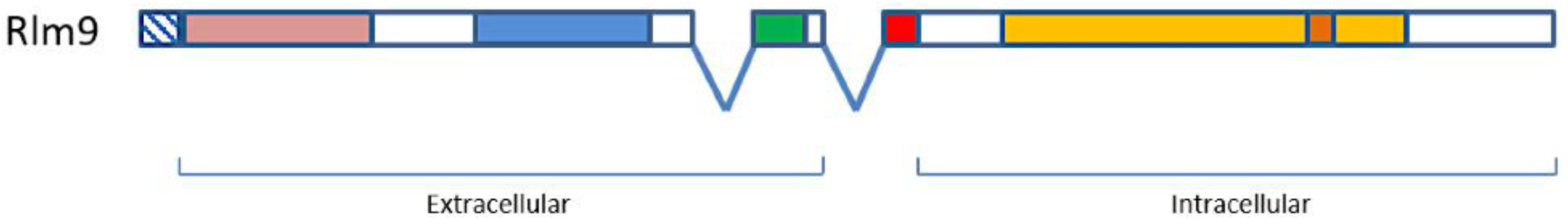
Domain organisation of the Rlm9 protein. The protein consists of 3 exons (introns denoted by ‘V’) and contains predicted signal peptide (hashed box), extracellular GUB_WAK pectin binding (light red), C-terminal WAK (blue) and EGF-like Ca2+ (green) domains, a transmembrane motif (red), and an intracellular serine/threonine protein kinase domain (light orange) with a guanylyl cyclase centre (dark orange).

BnaA07g20220D and its promoter was isolated from ‘Darmor’ and transferred to the susceptible *B. napus* cultivar ‘Westar N-o-1’. After transgenic events were analysed in the T_0_ generation by ddPCR, four independent transgenic lines, carrying between 1 and 9 copies of the transgene, were selected for phenotypic analysis. Self seed of each line (T_1_) was inoculated with the transgenic Lm isolate 2367:*AvrLm5-9.* Race specific resistance response was expressed in all Westar:BnaA07g20220D transgenic lines, confirming that the WAKL gene is indeed *Rlm9* (Figure 2). Further ddPCR analysis of the T_1_ plants derived from transgenic line NLA68 (one heterozygous transgene insertion at T_0_) allowed for selection of a T_1_ plant carrying a single homozygous insertion which was self-fertilized to produce homozygous T_2_ seed (hereafter referred to as Westar:*Rlm9*). Further testing of Westar:*Rlm9* with additional transgenic isolates carrying virulence genes matching other A07 blackleg *R* genes (2367:*AvrLm1*, 2367:*AvrLm3*, 2367:*AvrLm4-7*, 2367:*AvrLm7*) produced only susceptible interactions (data not shown) as previously demonstrated (Ghanbarnia *et al*., 2018). This reconfirmed both the identity of BnaA07g20220D as *Rlm9* and the specificity of the *Rlm9* – *AvrLm5-9* interaction.

**Figure 2.**
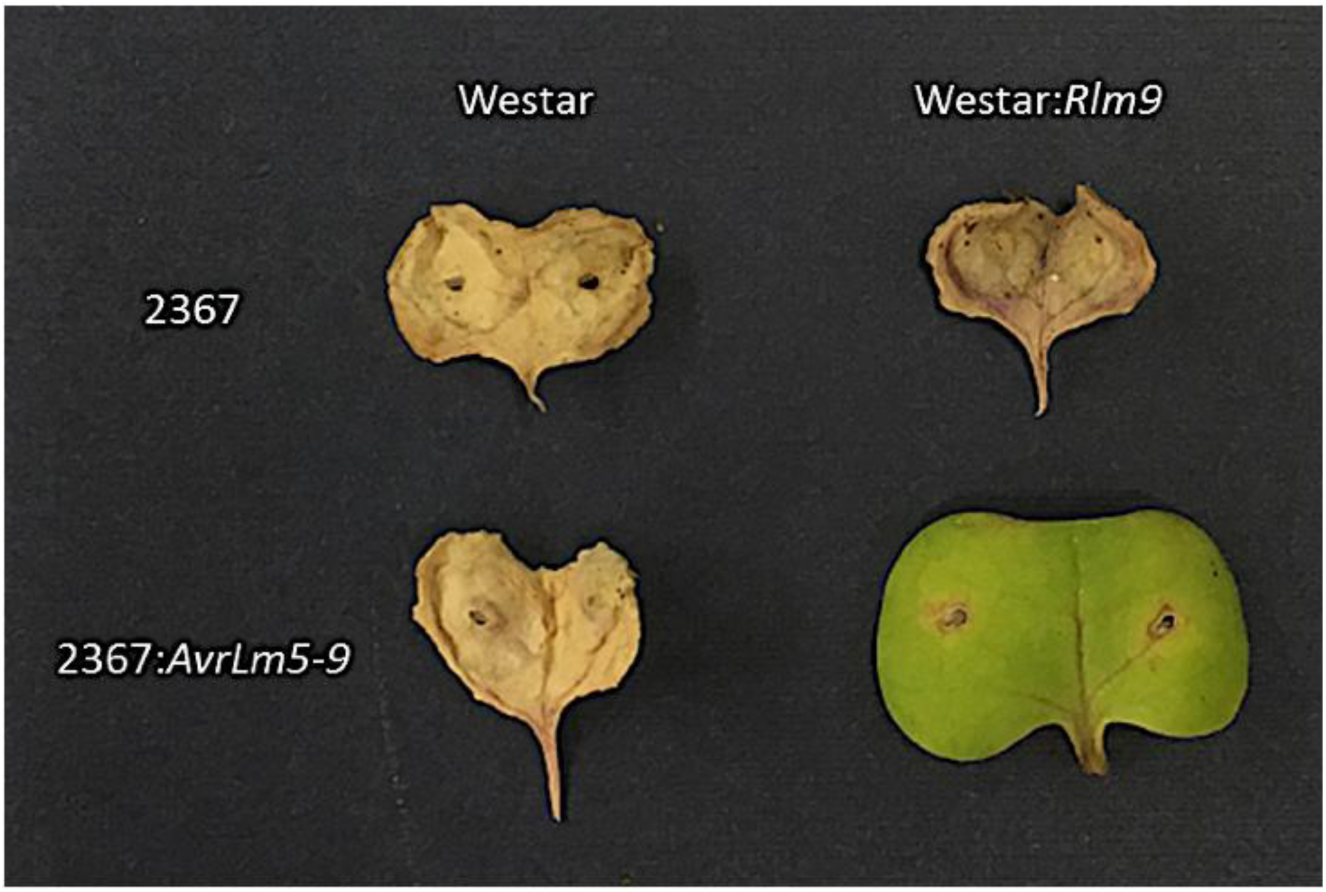
Transgenic Complementation of *Rlm9* Phenotype in *B. napus*. Cotyledons of Westar (no *R* gene) and Westar:*Rlm9* transgenic line infected with *L. maculans*, 14 days-post infection. Isolate 2367 (phenotype a9 – virulent towards *Rlm9*) and the transgenic isolate 2367:*AvrLm5-9* (phenotype A9 – avirulent toward *Rlm9*).

An identical *Rlm9* allele was identified from the genome sequence of *B. napus* var. ‘Tapidor’ (Bayer *et al*., 2017), which also harbours *Rlm9* (Larkan *et al*., 2016) (Supplementary Table 1). A susceptible allele (*rlm9*; Supplementary Table 1) was obtained from the genome sequence (v2.0) of *B. napus* var. ‘ZS11’ (He *et al*., 2015). The *B. rapa* var. ‘Chiifu’ homologue (Bra003598 – unknown *Rlm9* phenotype) (Wang et al., 2011) was also include for comparison studies. Comparison of the resistant Rlm9 and susceptible rlm9 proteins (95.72% identity overall) revealed that most of the variation appeared concentrated in the predicted pectin-binding (GUB_WAK) domains (15 substitutions within the 119 aa domain; 87.39% identity), while the C-terminal WAK and EGF-like domains were well conserved (94.69% and 100% identity, respectively).

RNAseq time course analysis revealed a significant upregulation of *Rlm9* during *L. maculans* infection (isolate 00-100; A2-3-5-6-(8)-9-10-L1-L2-L4) with approximately 5-fold increase early in the infection (3 days post-infection, FDR <0.03) and a near 8-fold increase in transcript abundance detected at 6 dpi (FDR<0.001) in the *Rlm9* variety ‘Darmor’ when compared to a susceptible (*rlm9*) variety ‘Topas DH16516’ or the mock (water) inoculated control. No significant difference was detected in the expression of the fungal *AvrLm5-9* between the inoculated susceptible and resistant lines over the same time course (Figure 3). Expression of the *B. napus SOBIR1* and *BAK1* genes, previously shown to interact with the *B. napus* RLP-type *R* genes, *Rlm2* and *LepR3* (Larkan *et al*., 2013; Larkan *et al*., 2015), were up-regulated (up to 10.5 fold) during infection of the *Rlm2* plants. However, in the infected *Rlm9* plants, both the *SOBIR1* and *BAK1* homologues (6 copies each in *B. napus*) showed little upregulation (0-2 fold), suggesting they may not be involved in the same manner during the WAKL *R*-gene response (Supplementary Figure 1).

**Figure 3.**
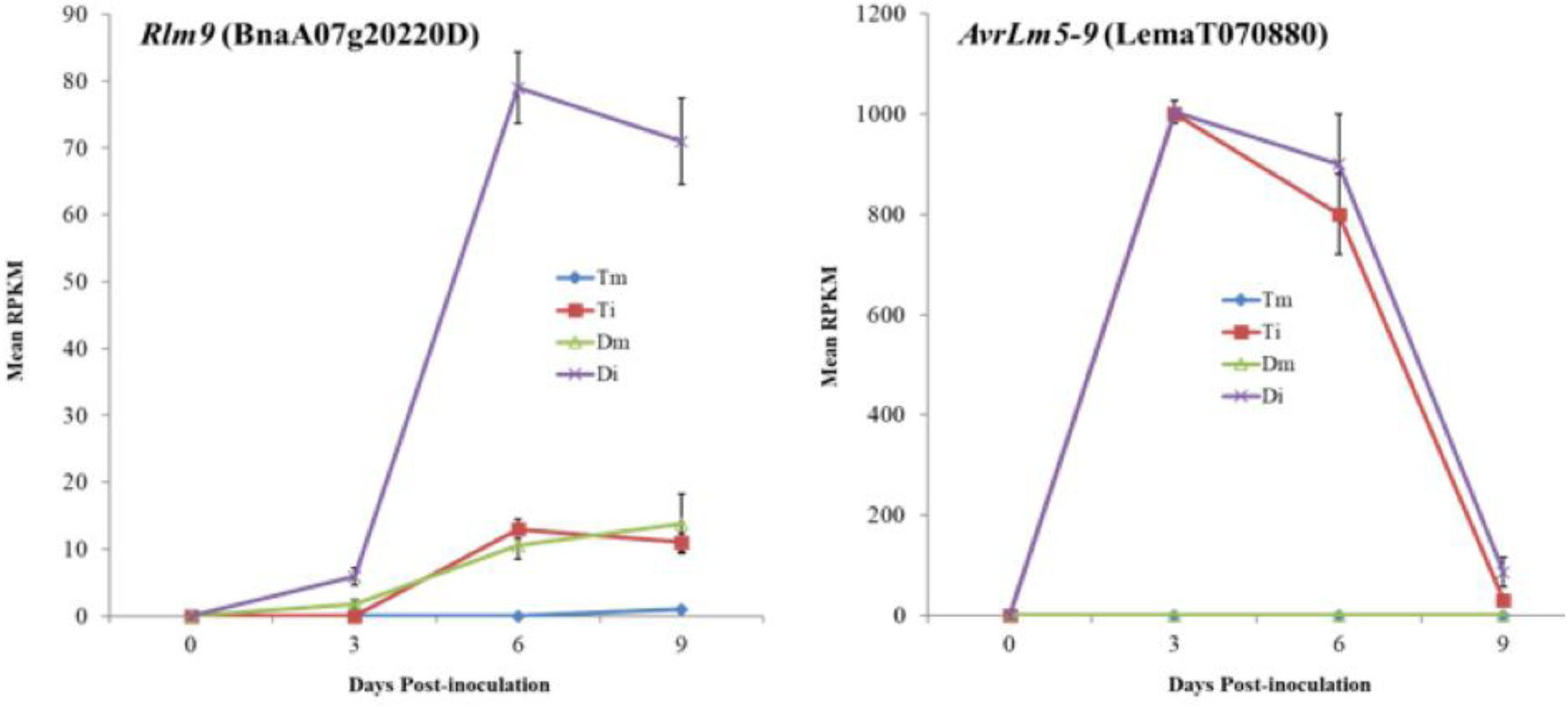
Expression Profile of *Rlm9* and *AvrLm5-9* Alleles during infection by *L. maculans* isolate 00-100. A) Mean RPKM (Reads Per Kilobase of transcript per Million mapped reads) for mock (m) and *L. maculans* infected (i) cotyledon lesions from *B.napus* lines Topas DH16516 (T – *rlm9*) and Darmor (D – *Rlm9*), showing significant upregulation of *Rlm9* detected between both Dm and Di, and Di and Ti, at all timepoints after zero. B) Mean RPKM for fungal *AvrLm5-9* during the same experiment, showing no significant difference between expression between Di and Ti.

### Confirmation of *Rlm9* in *B. napus* varieties

A selection of 22 *B. napus* cultivars, including many either previously identified as *Rlm9* lines, or suspected to harbour *Rlm9* based on previous differential pathology (data not shown), and the introgression line Topas-*Rlm9*, were tested for the presence of the *Rlm9* allele. The presence of *Rlm9* was first confirmed via infection with the transgenic *L. maculans* isolate 2367:*AvrLm5-9*, which induced a hypersensitive response in all 13 *Rlm9* lines (Supplementary Table 1, Supplementary Figure 2). All lines were susceptible to the non-transgenic 2367 isolate. The allele was successfully amplified from each of the 13 *Rlm9* lines (Supplementary Table 1), while only weak non-specific amplicons were produced from non-*Rlm9* lines, including cultivars containing other A07 blackleg *R* genes (*Rlm1, Rlm3, Rlm4* & *Rlm7*).

### No direct physical interaction detected between Rlm9 and either AvrLm5-9 or AvrLm4-7

Recently, we reported the cloning of *L. maculans* effector *AvrLm5-9* (Ghanbarnia *et al*., 2018). As previously reported recognition of AvrLm5-9 and AvrLm3 by their cognate R proteins, Rlm9 and Rlm3 is masked in the presence of AvrLm4-7 and this masking effect is neither due to direct interaction between these effector proteins nor due to the suppression of their transcription (Plissonneau *et al*., 2016; Ghanbarnia *et al*., 2018). To examine whether AvrLm5-9 directly interacts with Rlm9, we cloned the extracellular region of *Rlm9* in the prey vector pGADT7 and *AvrLm5-9* lacking the signal peptide sequences in the bait vector pGBKT7 for yeast two-hybrid assay. The assay was performed by co-transforming the bait and prey constructs to yeast strain Y2HGold. The combination of the *L. maculans* effector AvrLm1 (bait) and its *B. napus* host-interacting protein BnMPK9 (Ma *et al*., 2018) (prey) were used as a positive control. No interaction could be detected between AvrLm5-9 and the extracellular region of Rlm9 (Figure 4). To assess whether AvrLm4-7, which masks the recognition of AvrLm5-9 by Rlm9, directly interacts with either the extracellular region or the kinase domain of Rlm9 to supress *Rlm9*-mediated resistance, we co-transferred the bait vector pGBKT7:*ΔAvrLm4-7* and either prey vector pGADT7:*Rlm9*-EX or pGADT7:*Rlm9*-KD to yeast. As shown in Figure 4 there was no interaction between AvrLm4-7 and Rlm9-EX or Rlm9-KD, indicating that the masking of *Rlm9*-mediated resistance by AvrLm4-7 is not due to the direct interaction of AvrLm4-7 and Rlm9.

**Figure 4.**
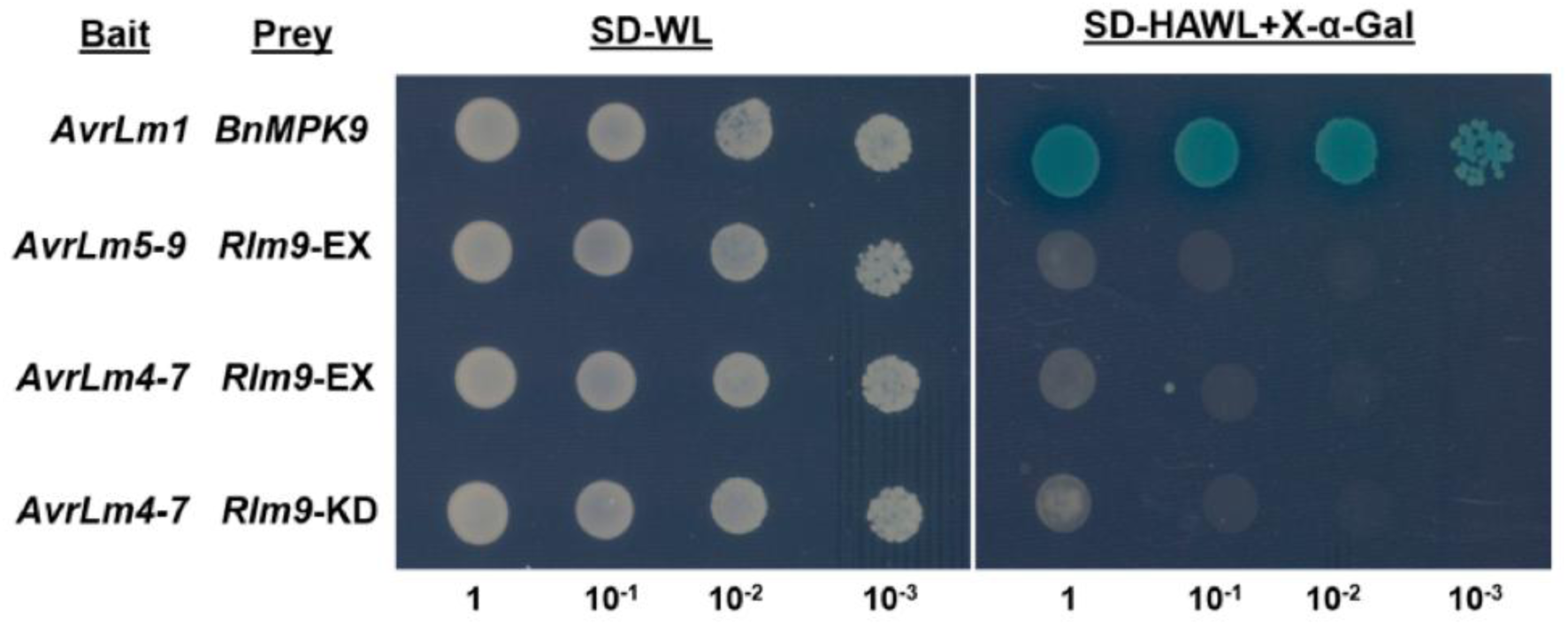
Yeast Two-hybrid Assay. Co-transformed (bait + prey) yeast plated at four dilutions (1 to 10-3). Interaction indicated by blue colouring. No interaction was detected for either AvrLm5-9 or AvrLm4-7 with the extracellular domain of Rlm9 (*Rlm9*-EX), nor AvrLm4-7 with the intracellular kinase domain (*Rlm9*-KD). AvrLm1-BnMPK9 was included as positive control.

### Evolution of the WAKL gene family in *B. napus*

To compare the evolution of the WAKL gene family between *A. thaliana* and *B. napus*, predicted WAKL-encoding genes, those encoding proteins with homology to the external domain of Rlm9, were extracted from the Darmor-*bzh* reference *B. napus* genome. After annotation 18 additional genes encoding potential functional RLKs (predicted proteins containing SP, TM and PK domains) were identified. All 18 predicted proteins also contained GUB_WAK and C-terminal WAK domains, while 15 of the 18 contained EGF-Like domains. WAKLs were distinguished from WAKs based on homology (>25% identity to extracellular domain of Rlm9), presence of C-terminal WAK domain and lack of twin EGF domains found in WAKs (Supplementary Table 2). The predicted *B. napus* WAKLs were found to be largely clustered on two chromosomes in each of the A (A08 & A09) and C (C06 & C08) genomes (Figure 5). Genomic alignment between syntenic *A. thaliana* blocks containing 18 of the previously-characterised 22 AtWAKLs (Verica & He, 2002) which also encode intact RLKs, and the two genomes of *B. napus* revealed that almost all of the predicted WAKLs in *B. napus* are syntenic to the WAKL genes clustered on *A. thaliana* chromosome 1 (Figure 5). Comparison of WAKL GUB_WAK domains to AtWAK1 showed that none of the four amino acid residues previously identified as contributing to homogalacturonan binding within the AtWAK1 GUB_WAK domain (Decreux *et al*., 2006) are conserved in either the resistant or susceptible alleles of Rlm9, nor in AtWAKL10, and show generally poor conservation in both WAKL and WAK predicted proteins from *B. napus*. (Supplementary Figure 3). Phylogenetic analysis for the predicted GUB_WAK domains also suggests that the WAKL proteins are a distinct evolutionary group from WAKs (Supplementary Figure 3.)

**Figure 5.**
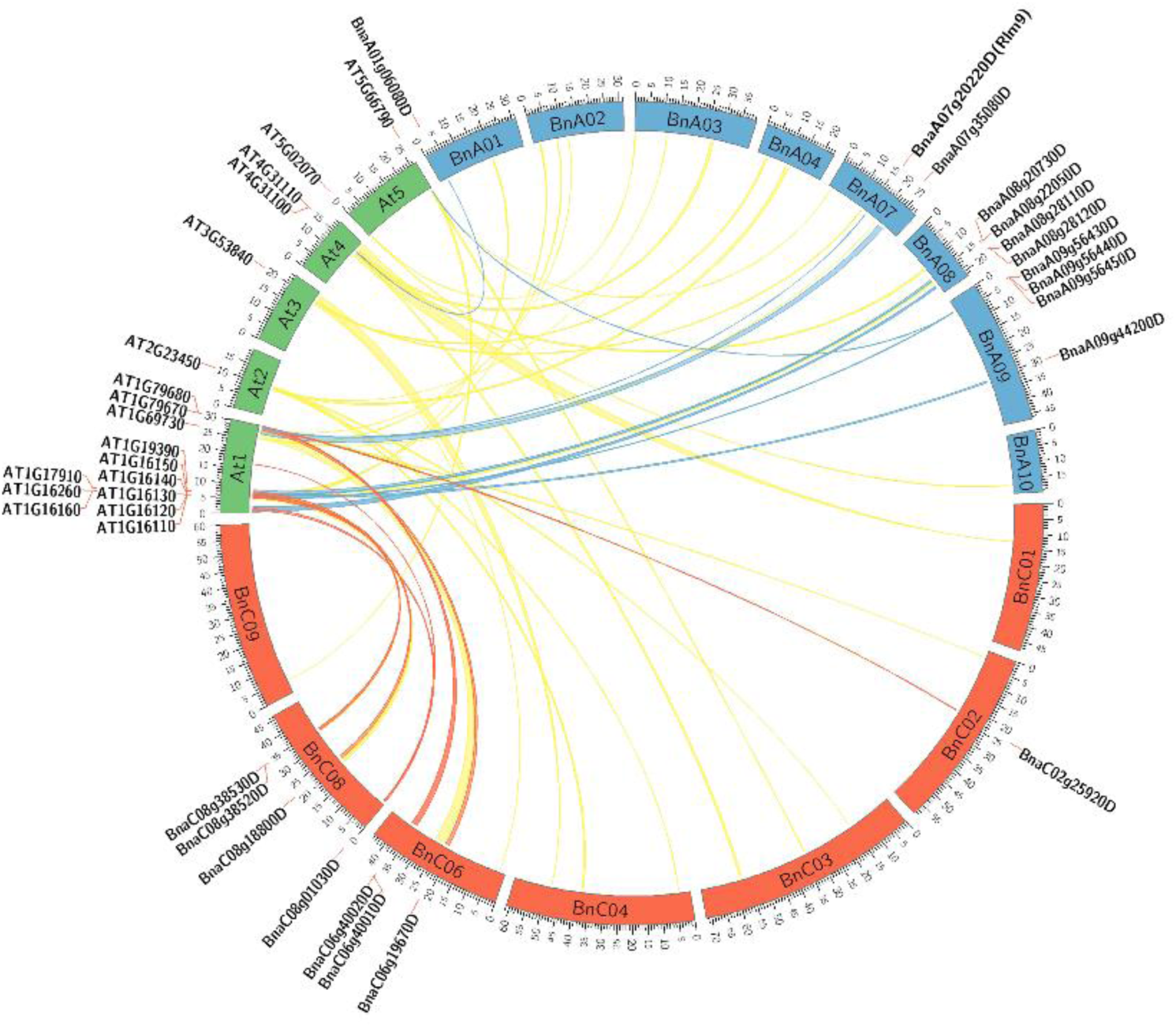
Syntenic alignment of *A. thaliana* and *B. napus* WAKL genomic regions. Genomic alignment between *A. thaliana* genomic blocks containing WAKL genes (“AT…” labels) and their syntenic matches in the *B. napus* A and C genomes. *B. napus* genes predicted to encode intact WAKL proteins labeled “Bna…”. Syntenic links between *A. thaliana* and *B. napus* WAKLs indicated by blue (A genome) and red (C genome) ribbons. Yellow ribbons indicate syntenic matches where no corresponding *B. napus* WAKL was found.

## Discussion

The number of characterised race-specific resistance (*R*) genes has significantly expanded since the cloning of the first *R* gene in 1992, with the majority of described *R* genes encoding intracellular Nod-like receptors (NLRs) (Zhang *et al*., 2017; Kourelis & Van Der Hoorn, 2018). However, a number of cell surface receptor proteins, collectively referred to as plant pattern recognition receptors (PRRs) which are involved in the recognition of extracellular plant pathogens, both through PAMPs and specific effectors, have also been identified (Boutrot & Zipfel, 2017). *Rlm9* and the recently-cloned wheat *Stb6* are the only examples of a race-specific WAKL-type *R* genes to be reported to date. *Stb6* confers resistance to races of the apoplastic fungal pathogen *Zymoseptoria tritici* which produce the matching effector AvrStb6, though resistance is semi-dominant and conferred without a typical hypersensitive response (HR) (Zhong *et al*., 2017; Saintenac *et al*., 2018). In contrast *Rlm9* induces a clear, dominant HR at the site of infection, responding to the presence of the *L. maculans* avirulence protein AvrLm5-9 (Ghanbarnia *et al*., 2018). The emergence of WAKL proteins as new players in race-specific resistance brings with it many fundamental questions. Using Y2H we did not detect a direct interaction between AvrLm5-9 and Rlm9, which was also reported to be the case between Stb6 and AvrStb6 (Saintenac *et al*., 2018). Although Y2H is not an optimal test to detect direct interaction of membrane proteins it is possible that Rlm9 recognition of AvrLm5-9 is indirect and mediated by a host target molecule. One such molecule could be a DAMP such as pectin monomers. However, the mechanism by which these predicted pectin-binding proteins function as mediators of race specificity has yet to be determined. The Stb6 protein contains a predicted extracellular galacturonan-binding (GUB_WAK) domain but does not contain either the C-terminal WAK or EGF-like Ca^2+^ domains found in Rlm9 and most other WAKL proteins (Saintenac *et al*., 2018), which suggests that these domains could be dispensable for WAKL-mediated effector-triggered immune response. The concentration of variation in the GUB_WAK domain between the resistant Rlm9 and susceptible rlm9 proteins in comparison to the other well-conserved extracellular domains (C-terminal WAK and EGF-like domains) also suggests that the GUB_WAK domain may play a pivotal role in recognition of AvrLm5-9. While the GUB_WAK domain of the *A. thaliana* wall-associated kinase 1 (WAK1) protein has been demonstrated to bind cell-wall pectins (Decreux & Messiaen, 2005), and WAKL proteins have been suggested to be associated with the cell wall (Verica & He, 2002; Hou *et al*., 2005), the same pectin-binding activity has yet to be shown for the predicted GUB_WAK domains of the WAKL proteins and much research needs to be undertaken before we can determine how these RLKs function. However, at present it should not be assumed that these proteins retain the ability to bind pectin. It may be that the original function of the gene was as a general DAMP receptor, as for AtWAK1 (Brutus *et al*., 2010), and that these proteins later evolved into a more specialised role in the detection of proteinaceous ligands, in this case AvrLm5-9. As the conservation between WAKs and WAKLs appears in the EGF and kinase domains, rather than the putative pectin-binging regions, it may be more appropriate to consider WAKs and WAKLs as subsets of the EGF protein superfamily, rather than grouping both WAKs and WAKLs together as “wall-associated” proteins (Kohorn, 2016).

In *A. thaliana*, Rlm9 shares the highest protein homology with WAKL10 (At1g79680.1). *AtWAKL10* is co-expressed with several pathogen response genes during biotic interactions. The protein kinase domain has been shown to be a twin-domain, also having guanylyl cyclase (GC) activity (Meier et al., 2010). Rlm9 contains an identical GC motif (SFGVVLAELITGEK) within the PK domain. GCs convert guanosine 5’-triphosphate (GTP) into guanosine 3’,5’-cyclic monophosphate (cGMP), an important signaling molecule during biotic interactions (Durner *et al*., 1998; Gehring & Turek, 2017). Plant cGMP-binding proteins include several actors in defense response pathways, including hydrogen peroxide production (Donaldson *et al*., 2016). The potential GC activity of Rlm9 could be a key component of the hypersensitive response to *L. maculans* infection. Interestingly, the wheat Stb6 protein, which does not trigger a hypersensitive response (Saintenac et al., 2018), appears to lack a GC centre in its kinase domain.

A search for WAKL homologues within the *B. napus* genome showed far fewer intact genes than was expected. *B. napus* is an amphidiploid hybrid of *B. rapa* (A genome) and *B. oleracea* (C genome), with each diploid genome having evolved from a hexaploid ancestor, with some gene loss occurring over time (Parkin *et al*., 2005; Ziolkowski *et al*., 2006). Therefore, for each single *A. thaliana* gene there is generally expected to be six homologues within *B. napus* (Grant *et al*., 1998). Despite there being 22 WAKLs characterised in *A. thaliana* (Verica & He, 2002) we were only able to identify 29 intact WAKL genes predicted within the *B. napus* ‘Darmor’ sequence, only 18 of which are predicted to retain the SP, TM and PK domains required to function as an RLK (Supplementary Table 2). This suggests a disproportionate evolution of the genes since the Arabidopsis-Brassica split 20-24 Mya (Ziolkowski *et al*., 2006). This may be due to functional redundancy, making many of the WAKL homologues within the amphidiploid genome (AACC) of *B. napus* dispensable. The relative abundance of WAKL genes in *A. thaliana* may also be due to a higher rate of gene expansion, as evidenced by the dense clusters of WAKL homologues and abundant tandem duplications found on *A. thaliana* chromosome 1 (12 genes total -Verica & He, 2002) which are only partially represented within each of the *B. napus* A and C genomes as homoeologues, while several AtWAKLs found on other chromosomes do not appear to be represented by intact genes in *B. napus*, despite syntenic links between the genomes (Figure 5).

As we recently reported, the recognition of AvrLm5-9 by Rlm9 is masked in the presence of AvrLm4-7 (Ghanbarnia *et al*., 2018). Similarly, AvrLm3 recognition by Rlm3 is also masked in the presence of AvrLm4-7 (Plissonneau *et al*., 2016). With the cloning of *Rlm9* and the characterisation of the gene as a WAKL we now have the basis for possibly identifying the other three blackleg *R* genes within the *Rlm3*/*4*/*7*/*9* cluster, co-located on chromosome A07 (Larkan *et al*., 2016). Though *Rlm9* is the only A07 blackleg *R* gene carried by any of the published *B. napus* genomes (we identified the same *Rlm9* allele in both ‘Darmor’ (Chalhoub *et al*., 2014) and ‘Tapidor’ (Bayer *et al*., 2017), and showed a lack of specific *R* genes in ‘ZS11’ (He *et al*., 2015) during this study) previous investigations into the blackleg resistance carried by ‘DH12075’, a doubled-haploid F_1_-derived line from ‘Westar’ (no *R* gene) and ‘Cresor’ (*Rlm3* -Larkan *et al*., 2016) parents, identified three WAKL genes in the same location (I. Parkin, unpublished). We are currently investigating the allelic variation of the *Rlm9* locus in multiple *B. napus* accessions carrying *Rlm3, Rlm4* and *Rlm7* through parental-specific genome resequencing. If these genes prove to be variants or duplications of the same WAKL locus we can potentially gain further insight as to the evolution of WAKL-type *R* genes, and the *Brassica-Leptosphaeria* pathosystem may prove to be a model system by which the mechanism of fungal avirulence protein recognition by WAKLs can be determined.

## Methods

### Candidate Identification and Transformation

The BnaA07g20220D gene, including 1000 bp upstream and 500 bp downstream of the predicted CDS (4141 bp total), was PCR amplified (Q5 High-Fidelity 2x Master Mix, New England BioLabs) from ‘Darmor’ DNA, verified by Sanger sequencing, and transferred to the Gateway-compatible transformation vector pMDC123 (Curtis & Grossniklaus, 2003). The same primers (GW-DarWAKL F + R; Supplementary Table 3) were used to survey a selection of *B. napus* cultivars to confirm the presence of the target allele in multiple *Rlm9* sources. The genomic candidate construct was transformed via *Agrobacterium* into the susceptible *B. napus* cultivar ‘Westar N-o-1’ as previously described (Larkan *et al*., 2013). Homozygous, single-insertion transgenic plants were selected in the T_1_ generation using ddPCR (Larkan *et al*., 2015). Final confirmation of phenotype was performed using the transgenic *L. maculans* isolate 2367:*AvrLm5-9* (Ghanbarnia *et al*., 2018).

### Transcript Analysis

Transcription time course profiles were generated by RNA-Seq (Illumina) for the *B. napus* cultivars Topas DH16516 (no *R* gene) and Darmor (*Rlm9*) during infection with the *L. maculans* isolate 00-100 (avirulence profile A2-3-5-6-(8)-9-(10)-L1-L2-L4 (Larkan *et al*., 2016)) with sampling at 0, 3, 6 and 9 days after inoculation (dai). Additional data for the lines Topas-*Rlm2* and Topas-*Rlm3*, growth conditions, tissue sampling, RNA processing and read mapping protocols were as previously described (Becker *et al*., 2019; Haddadi *et al*., 2019). Confirmation of the predicted coding region and protein sequence was obtained by aligning merged RNA sequencing reads to the Darmor genome sequence using Bowtie2 (http://bowtie-bio.sourceforge.net/bowtie2/manual.shtml) and CLC Genomics Workbench 11 (https://www.qiagenbioinformatics.com/products/clc-genomics-workbench/).

### Yeast Two-hybrid Assay

For yeast two-hybrid assay, *AvrLm5-9* lacking signal peptide sequence was cloned into pGBKT7 bait vector with primer set Δsp*AvrLm5-9*-NcoI/Δsp*AvrLm5-9*-EcoRI and *AvrLm4-7* lacking signal peptide sequence was coned into pGBKT7 bait vector with primer set Δsp*AvrLm4*-*7*NdeI/Δsp*AvrLm4*-*7*-PstI. The intracellular kinase domain of *Rlm9* was cloned into pGADT7 prey vector with primer set *Rlm9*-KD-NdeI/*Rlm9*-KD-EcoRI and the extracellular region of *Rlm9* was cloned into pGADT7 with primer set *Rlm9*-EX-NedI/*Rlm9*-EX-EcoRI (Clontech, Mountain View, USA). We used the matchmaker GLA4 two-hybrid system and yeast strain Y2HGold (Clontech, Mountain View, USA). The yeast strain Y2HGold was co-transformed with bait and prey plasmid combinations using lithium-acetate and polyethylene glycol 3350 followed the manufacture manual. Transformants harboring both bait and prey plasmids were selected on plates containing minimal medium lacking Leu and Trp (SD-WL). Empty prey vector pGBKT7 or pGADT7 used as bait or prey served as controls. pGBKT7:Δ*AvrLm1* and pGADT7:*BnMPK9* were used as positive control (Ma *et al*., 2018). One colony per combination was picked from SD-WL plates to inoculate 1 mL SD-WL culture. After 36 h growth, cells were collected by centrifugation and resuspended in 25 µL 0.9% NaCl from OD_600_=1 to OD_600_=0.00001 and spotted on SD-WL and SD-AHWL plates supplementing with 40 μg/mL X-α-Gal (Clontech, Mountain View, USA) and 200 ng/ml Aureobasidin A (Clontech, Mountain View, USA). After 3 days incubation, the plates were checked for growth and photographed.

### Genomic and Phylogenetic Analyses

Predicted *B. napus* WAKL protein sequences, matching the extracellular domain of *Rlm9*, were retrieved from the Darmor-*bzh* reference annotation (Chalhoub *et al*., 2014) using the blastP function (default values) in CLC Genomics Workbench v12. Protein sequences were annotated using InterPro (http://www.ebi.ac.uk/interpro/). Only those proteins containing predicted domains required for RLK function (SP, TM & PK) were included as potential functional WAKLs (Supplementary Table 2). Visualization of the homology between *Arabidopsis thaliana* TAIR10 (Wensel *et al*., 2011) and *Brassica napus* Darmor-*bzh* (Chalhoub *et al*., 2014) was produced using Circos (Krzywinski *et al*., 2009). Orthologous gene pairs identified from synteny analysis were used to determine regions of homology and intersections with the defined *B. napus* WAKL genes using BEDTools intersect (Quinlan & Hall, 2010). Multiple sequence alignments and dendrograms were produced using CLC Workbench v7.9.1 software.

## Supporting information

main figures

## Acknowledgments

We would like to thank Elena Beynon, Colin Kindrachuk, Gordon Gropp, Catherine Guenther, Justin Skurdal and Erica Boyer for their technical assistance and Delwin Epp for generating the *B. napus* transgenic lines. Funding support for this project was provided by SaskCanola and Agriculture and Agri-Food Canada.

## Authors’ contributions

NJL and MHB conceived the study. NJL designed the experiments and performed the research together with LM, PH and MD. MB and PH performed the bioinformatics analysis. NJL, MHB and IAPP wrote the manuscript with technical contributions from LM, PH and MB. All authors have read and approved the current version of the manuscript.

---

**Supplementary Figure 1.Expression of *B. napus SOBIR1* and *BAK1* Homologues.** Expression of genes during infection by *L. maculans* relative to Topas DH16516 (no *R* gene) in *B. napus* lines carrying the *R* genes *Rlm9* (WAKL), *Rlm3* (suspected WAKL) or *Rlm2* (RLP).

**Supplementary Figure 2. Transgenic Confirmation of *Rlm9* Phenotype in *B. napus* Varieties.** Cotyledons of *Rlm9 B. napus* cultivars, 14 days after infection.

**Supplementary Figure 3. Multiple Sequence Alignment and Dendrogram for GUB_WAK Domains.** A) Alignment and B) dendrogram of GUB_WAK domains predicted for *B. napus* WAKLs and WAKs with >20% amino acid identity to the extracellular domain of Rlm9. Black boxes (A) indicate residues previously identified as contributing to pectin binding in AtWAK1.

**Supplementary Table 1. *Rlm9* PCR and Pathological Interactions for *B. napus* Lines.** Isolate interactions classified as either virulent (avr) or avirulent (Avr) based in median infection score (in brackets, 0-9 scale – Larkan et al., 2013).

**Supplementary Table 2. *B. napus* WAKL and WAK protein matches**

**Supplementary Table 3. PCR Primers**

