## Supplementary material for "The *Brassica napus* Wall-Associated Kinase-Like (WAKL) gene *Rlm9* provides race-specific blackleg resistance": main figures

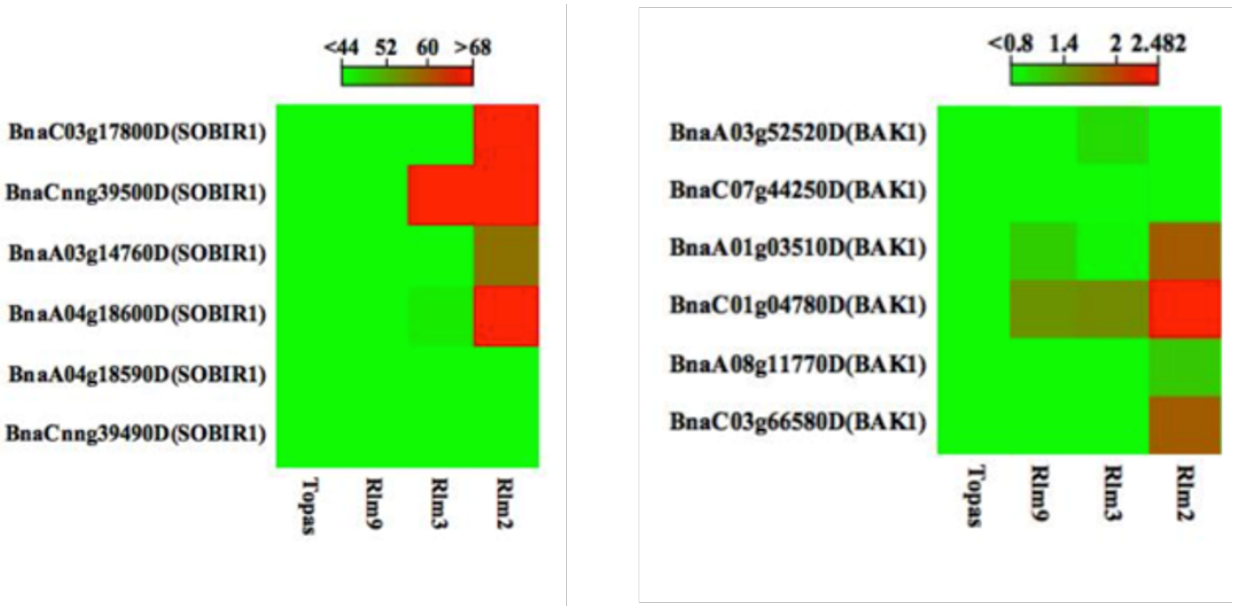

**Supplementary Figure 1. Expression of *B. napus SOBIR1* and *BAK1* Homologues.** Expression of genes during infection by *L. maculans* relative to Topas DH16516 (no *R* gene) in *B. napus* lines carrying the *R* genes *Rlm9* (WAKL), *Rlm3* (suspected WAKL) or *Rlm2* (RLP).

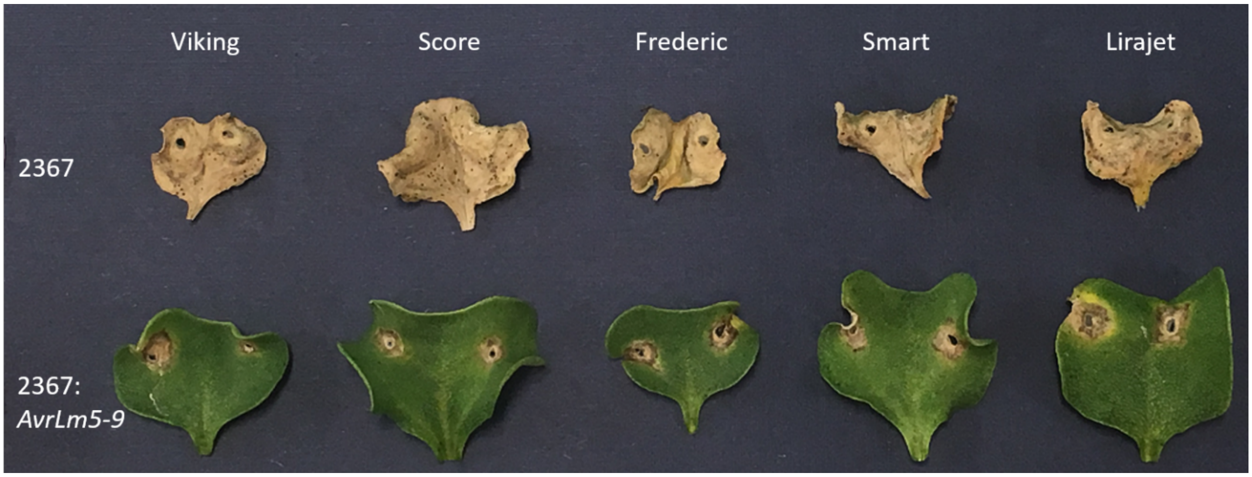

**Supplementary Figure 2. Transgenic Confirmation of *Rlm9* Phenotype in *B. napus* Varieties.** Cotyledons of *Rlm9 B. napus* cultivars, 14 days after infection.

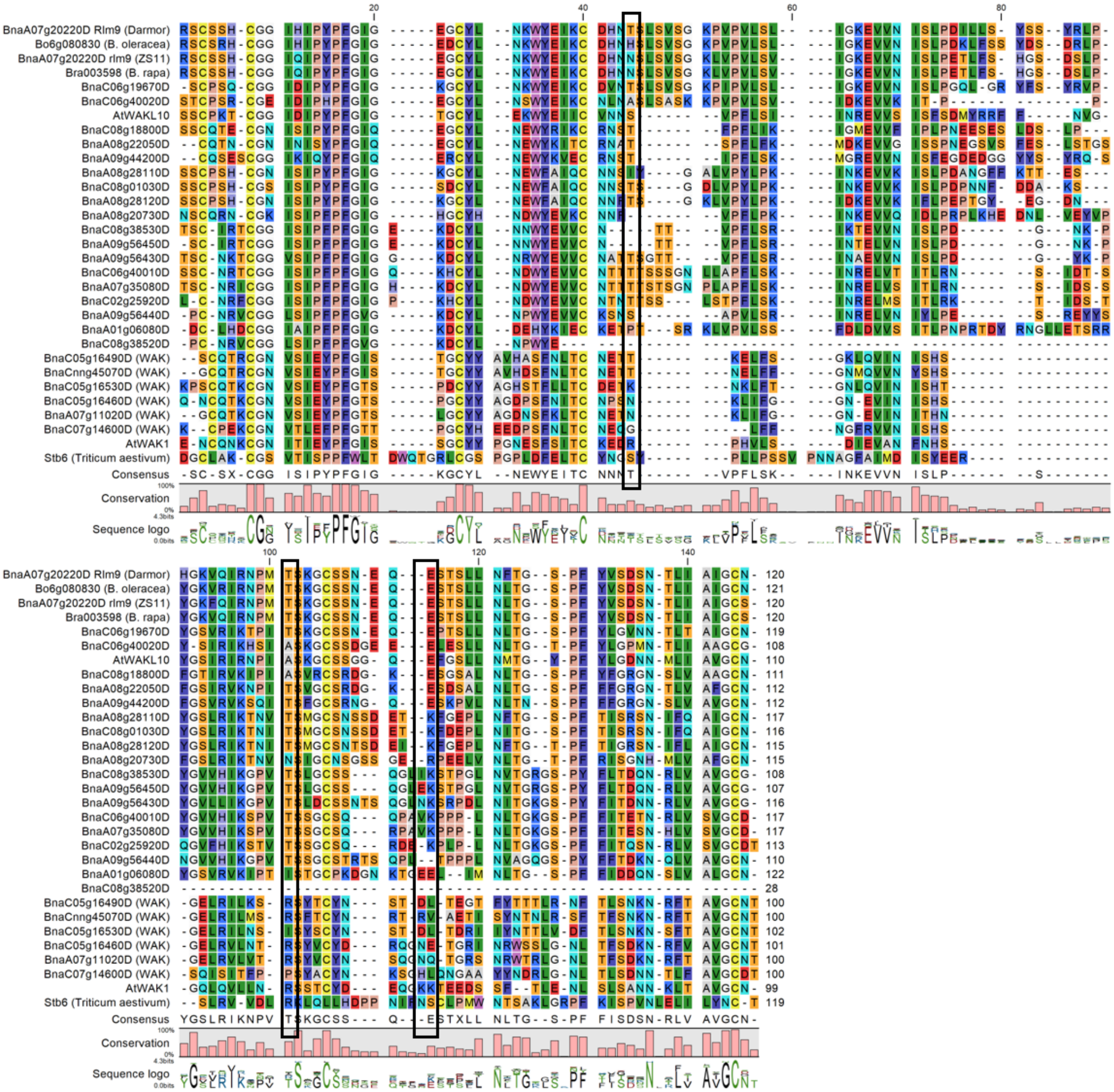
BnaA07g20220D Rlm9 (Darmor)
Bo6g080830 (B. oleracea)
BnaA07g20220D rlm9 (ZS11)
Bra003598 (B. rapa)
BnaC06g19670D
BnaC06g40020D
AtWAKL10
BnaC08g18800D
BnaA08g22050D
BnaA09g44200D
BnaA08g28110D
BnaC08g01030D
BnaA08g28120D
BnaA08g20730D
BnaC08g38530D
BnaA09g56450D
BnaA09g56430D
BnaC06g40010D
BnaA07g35080D
BnaC02g25920D
BnaA09g56440D
BnaA01g06080D
BnaC08g38520D
BnaC05g16490D (WAK)
BnaCnng45070D (WAK)
BnaC05g16530D (WAK)
BnaC05g16460D (WAK)
BnaA07g11020D (WAK)
BnaC07g14600D (WAK)
AtWAK1
Stb6 (Triticum aestivum)
Consensus
Conservation
Sequence logo

A)

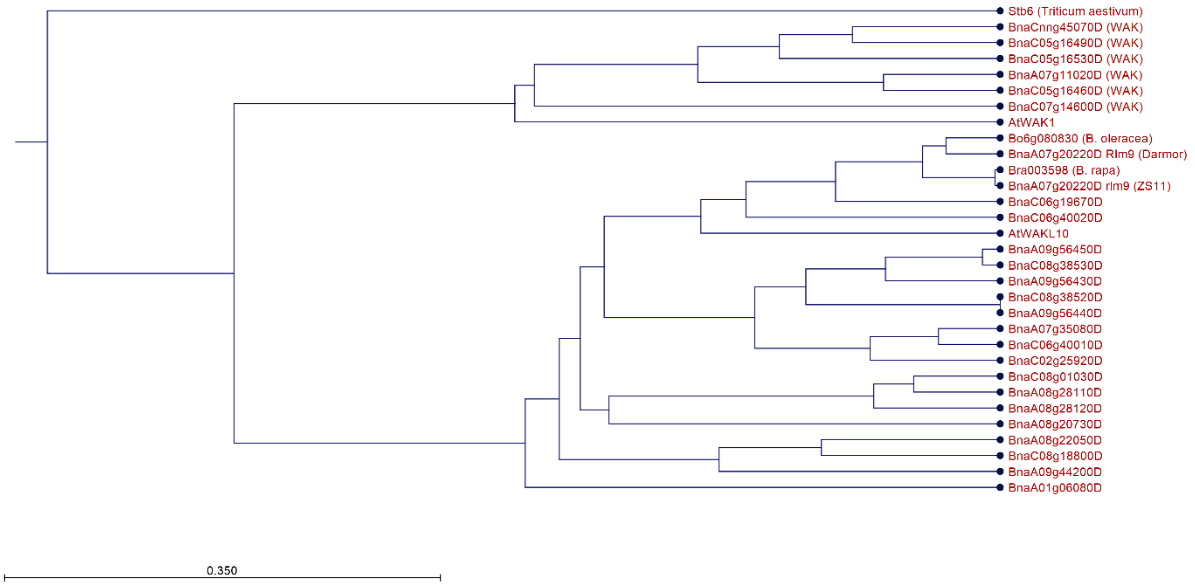

**B)**

**Supplementary Figure 3. Multiple Sequence Alignment and Dendrogram for GUB_WAK Domains.** A) Alignment and B) dendrogram of GUB_WAK domains predicted for *B. napus* WAKLs and WAKs with >20% amino acid identity to the extracellular domain of Rlm9. Black boxes (A) indicate residues previously identified as contributing to pectin binding in AtWAK1.

**Supplementary Table 1. *Rlm9* PCR and Pathological Interactions for *B. napus* Lines.** Isolate interactions classified as either virulent (avr) or avirulent (Avr) based in median infection score (in brackets, 0-9 scale [12]).

| *B. napus* Variety | *R* gene | *Rlm9* PCR | 2367s | 2367:*AvrLm9* |
|---|---|---|---|---|
| Topas DH16516 | none | - | avr (9) | avr (9) |
| Westar N-o-1 | none | - | avr (9) | avr (9) |
| ZS11 | none | - | avr (9) | avr (9) |
| Darmor | *Rlm9* | + | avr (9) | Avr (3) |
| Goeland | *Rlm9* | + | avr (9) | Avr (3) |
| DH12075 | *Rlm3* | - | avr (9) | avr (9) |
| Columbus | *Rlm1, Rlm3* | - | avr (9) | avr (9) |
| Quantum | *Rlm3* | - | avr (9) | avr (9) |
| Falcon | *Rlm4* | - | avr (9) | avr (9) |
| Scoop | *Rlm4* | - | avr (9) | avr (9) |
| Roxet | *Rlm7* | - | avr (9) | avr (9) |
| Bristol | *Rlm2, Rlm9* | + | avr (9) | Avr (3) |

| | | | | |
|---|---|---|---|---|
| Samurai | *Rlm2, Rlm9* | + | avr (8) | Avr (3) |
| Tapidor DH | *Rlm2, Rlm9* | + | avr (7) | Avr (3) |
| Frederic | unknown | + | avr (9) | Avr (4) |
| Lirajet | unknown | + | avr (7) | Avr (3) |
| Mohican | unknown | + | avr (9) | Avr (3) |
| Prince | unknown | + | avr (9) | Avr (4) |
| Score | unknown | + | avr (9) | Avr (3) |
| Smart | unknown | + | avr (9) | Avr (3) |
| Viking | unknown | + | avr (9) | Avr (3) |
| Wotan | unknown | + | avr (9) | Avr (3.5) |

**Supplementary Table 2. *B. napus* WAKL and WAK protein matches**

| | blastP Scores | | | | Predicted Domains | | | | | | |
|---|---|---|---|---|---|---|---|---|---|---|---|
| Gene | Max score | Min E-value | % Identity | | SP | GUB_WAK | WAK | EGF-Like | TM | PK | |
| BnaA07g20220D (*Rlm9*) | 1,942.00 | 0 | 100 | | y | y | y | y | y | y | |
| BnaC06g19670D | 1,443.00 | 0 | 77.45 | | y | y | y | y | y | y | |
| BnaC06g19690D | 1,185.00 | 5.95E-158 | 79.29 | | n | n | y | n | y | y | |
| BnaA07g35110D | 743 | 1.66E-95 | 50.69 | | y | y | y | y | y | n | |
| BnaC06g40020D | 758 | 2.43E-93 | 50 | | y | y | y | y | y | y | WAKLs |
| BnaA09g44200D | 640 | 2.51E-76 | 43.45 | | y | y | y | n | y | y | |
| BnaC08g18800D | 601 | 2.93E-71 | 41.37 | | y | y | y | y | y | y | |
| BnaA08g28120D | 606 | 3.18E-71 | 41 | | y | y | y | n | y | y | |
| BnaA08g28110D | 599 | 5.36E-71 | 40.33 | | y | y | y | y | y | y | |
| BnaA08g22050D | 583 | 7.77E-68 | 40.27 | | y | y | y | y | y | y | |
| BnaC08g01030D | 580 | 1.49E-67 | 40 | | y | y | y | y | y | y | |
| BnaC07g43130D | 553 | 4.84E-64 | 39.5 | | y | y | y | y | n | y | |
| BnaA01g06080D | 536 | 3.67E-61 | 38.83 | | y | y | y | y | y | y | |
| BnaA06g38500D | 512 | 1.26E-58 | 39.7 | | n | y | y | n | y | n | |
| BnaC08g38530D | 508 | 1.75E-57 | 39.83 | | y | y | y | y | y | y | |
| BnaA09g56450D | 511 | 5.37E-57 | 39.49 | | y | y | y | y | y | y | |
| BnaC01g07330D | 476 | 4.97E-56 | 38.23 | | y | y | y | n | n | n | |
| BnaC06g40010D | 493 | 3.07E-55 | 36.65 | | y | y | y | y | y | y | |
| BnaA09g56440D | 490 | 9.04E-55 | 34.46 | | y | y | y | y | y | y | |
| BnaA07g35080D | 482 | 1.69E-54 | 36.75 | | y | y | y | y | y | y | |
| BnaC02g25920D | 483 | 4.25E-54 | 34.86 | | y | y | y | y | y | y | |
| BnaA08g20730D | 477 | 4.04E-53 | 35.12 | | y | y | y | y | y | y | |
| BnaA09g56430D | 471 | 1.96E-52 | 37.64 | | y | y | y | n | y | y | |
| BnaC08g38510D | 417 | 1.51E-45 | 35.04 | | y | y | y | n | n | y | |
| BnaA08g23630D | 410 | 1.80E-44 | 32.95 | | y | y | y | y | n | y | |
| BnaA01g06050D | 391 | 1.89E-41 | 35.99 | | y | y | y | n | n | y | |
| BnaC05g49930D | 369 | 5.27E-40 | 31.58 | | n | y | y | n | y | n | |
| BnaC03g57020D | 366 | 5.41E-38 | 29.59 | | n | y | y | y | y | y | |
| BnaC01g07320D | 337 | 1.15E-34 | 32.37 | | n | y | y | y | n | y | |
| BnaC08g38520D | 312 | 1.21E-30 | 25.99 | | y | y | y | y | y | y | |
| BnaC05g16530D | 180 | 4.37E-14 | 24.36 | | y | y | n | 2x | y | n | |
| BnaC05g16490D | 181 | 6.15E-14 | 23.74 | | y | y | n | 2x | y | y | |
| BnaC05g16460D | 168 | 2.97E-12 | 26.72 | | y | y | n | 2x | y | y | |
| BnaCnng45070D | 164 | 6.19E-12 | 23.46 | | y | y | n | 2x | y | y | |
| BnaA07g11020D | 154 | 1.55E-10 | 24.86 | | y | y | n | 2x | y | y | WAKs |
| BnaC07g14600D | 150 | 4.64E-10 | 25.82 | | y | y | n | 2x | y | y | |
| BnaC07g14590D | 130 | 1.47E-07 | 24.77 | | y | y | n | n | n | y | |
| BnaC02g23840D | 125 | 5.22E-07 | 23.65 | | y | y | n | y | y | y | |
| BnaAnng15620D | 94 | 3.33E-03 | 22.35 | | y | y | n | 2x | y | y | |
| BnaA08g21390D | 94 | 3.56E-03 | 21.57 | | y | y | n | 2x | y | y | |

### Supplementary Table 3. PCR Primers

| | |
|---|---|
| GW-DarWAKL-F | GGGGACAAGTTTGTACAAAAAAGCAGGCTTCGCGGGAGACCTAGACTAGATCGACAA |
| GW-DarWAKL-R | GGGGACCACTTTGTACAAGAAAGCTGGGTCCACCATTACGAAGGCCACACGC |
| ΔspAvrLm47-NdeI | GCGGCATATGTGTAGAGAGGCCTCAATATCTGGAG |
| ΔspAvrLm47-PstI | GCGGCTGCAGTTAGTCGCAACCACGAGTCCTT |
| Rlm9-KD-NdeI | GCGGCATATGTTCAGTTTAACTAGAATACTTGGCCAAGG |
| Rlm9-KD-EcoRI | GCGGGAATTCTTAAATTTTCTCCAATTCCATGGACACTT |
| Rlm9-EX-NdeI | GCGGCATATGATGAGCTCTTATAATCTTTCTTCCCTTTTC |

| | |
|---|---|
| Rlm9-EX-EcoRI | GCGGGAATTCAGCAAGCCGGTGATTGCTATACAC |
| ΔspAvrLm5-9-NcoI | GCGGCCATGGAGCACGACTGCCATCAGGTCACC |
| ΔspAvrLm5-9-EcoRI | GCGGGAATTCCTAATGCTTTAGACAATGAACATTAGGATTAG |
| Rlm9-FB | ACAAGTTTGTACAAAAAAGCAGGCTTTATGAGCTCTTATAATCTTTCTTCCCTTTTC |
| Rlm9-RB-S | ACCACTTTGTACAAGAAAGCTGGGTCCCGTGTTTGGTAAGGAAACAGAGG |
